# Mode of Targeting to the Proteasome Determines GFP Fate

**DOI:** 10.1101/2020.07.15.205310

**Authors:** Christopher E. Bragança, Daniel A. Kraut

## Abstract

The Ubiquitin-proteasome system (UPS) is the canonical pathway for protein degradation in eukaryotic cells. Green fluorescent protein (GFP) is frequently used as a reporter in proteasomal degradation assays. However, there are multiple variants of GFP in use, and these variants have different stabilities. We previously found that the proteasome’s ability to unfold and degrade substrates is enhanced by polyubiquitin chains on the substrate, and that proteasomal ubiquitin receptors mediate this activation. Herein we investigate how the fate of GFP variants of differing stabilities is determined by the mode of targeting to the proteasome. We compared two targeting systems: linear Ub_4_ degrons and the UBL domain from yeast Rad23, both of which are commonly used in degradation experiments. Surprisingly, the UBL degron allows for degradation of the most stable sGFP-containing substrates, while the Ub_4_ degron does not. Destabilizing the GFP by circular permutation allows degradation with either targeting signal, indicating that domain stability and mode of targeting combine to determine substrate fate. Finally, we show that the ubiquitin receptor Rpn13 is primarily responsible for the enhanced ability of the proteasome to degrade stable UBL-tagged substrates.

---

The ubiquitin-proteasome system (UPS) is responsible for the bulk of protein degradation in eukaryotic cells. Proteins to be degraded are typically polyubiquitinated on one or more lysines by the action of E1, E2 and E3 enzymes. The polyubiquitin chain is recognized by one of three ubiquitin receptors on the 19S subunit of the proteasome, Rpn1, Rpn10 or Rpn13, or by ubiquitin shuttle proteins that bind to both polyubiquitin and one of the ubiquitin receptors. The substrate is then engaged by the ATP-dependent Rpt motor proteins at an unstructured initiation region and translocated into the 20S core particle’s central degradation chamber. The polyubiquitin chain passes by the deubiquitinase Rpn11 during translocation, which removes the chain, allowing ubiquitin to be recycled for additional rounds of protein targeting. Although recent advances have led to the understanding of many of the details of substrate recognition and unfolding, it remains unclear why three ubiquitin receptors (and multiple adaptors) are necessary or what controls whether a protein targeted to the proteasome by ubiquitination is successfully unfolded or escapes degradation.

Green fluorescent protein (GFP) and its derivatives are some of the most commonly used reporters used to study the UPS both *in vitro* and *in vivo* (1). For example, N-end rule (Ub-X-GFP) and ubiquitin fusion degradation (UFD; Ub^G76V^-GFP) substrates (2) have been used to explore the effects of Gly-Ala repeats, Huntingtin protein fragments and prions on global protein degradation (3–7), proteasome inhibitors in trypanosomes (8), and many other aspects of *in vivo* degradation. Other GFP-based systems have also been proposed for global monitoring of protein degradation in cells (9, 10). GFP is commonly fused to proteins of interest to determine whether and how fast they will be degraded in living cells, or even to control their degradation (e. g. (11–13)). GFP has also been used extensively in *in vitro* investigations of proteasome activity and mechanism (14–19).

However, there are challenges to using GFP as a model substrate. GFP forms an exceptionally stable 11-stranded ß-barrel. Extraction of a single ß strand from the barrel by ATP-dependent proteases leads to a still stable 10-stranded intermediate, and the original ß-barrel can fully reform if pulling is not rapid enough (20). Indeed, GFP-containing fragments as an endpoint of degradation have been observed both in cells and *in vitro* (16); these partial-degradation events are potentially frequently missed as most experiments simply look at total fluorescence. On the other hand, circular permutants of GFP that unfold in a single step have been shown to be degraded by the bacterial ATP-dependent protease ClpXP without stalling (20). The proteasome has been suggested to be even more processive than ClpXP (21) and thus capable of degrading even difficult-to-unfold substrates like GFP without stalling (17). However, in some cases GFP unfolding and degradation in cells requires the unfoldase cdc48/p97 in addition to the proteasome’s own unfoldase activity (22–24). Thus, a better understanding of GFP unfolding by the proteasome will inform the interpretation of a diversity of experiments.

We recently showed that substrate ubiquitination can increase the proteasome’s unfolding ability, and that proteasomal ubiquitin receptors mediate this increase, with Rpn13 playing the largest role (16, 25). Herein we set out to determine if the degradation of GFP is be affected by the mode of targeting to the proteasome, and, if so, which ubiquitin receptors are responsible. We used two previously established targeting systems, the linear Ub_4_ modification, with four non-hydrolyzable ubiquitins connected by short linkers, and the ubiquitin-like (UBL) domain from yeast Rad23, both attached to the N-terminus. In both cases, addition of an unstructured initiation region at the C-terminus of a target protein has been shown to lead to efficient degradation by the proteasome (26). Rpn1 has been suggested to be the primary receptor for Rad23 (27), while Rpn10 and Rpn13 are thought to be the primary receptors for ubiquitinated substrates (28). Quite recently, the Matouschek lab used a circular permutant of GFP to show that indeed Rpn10, and to a lesser extent Rpn13 were the only receptors used by Ub_4_, while Rpn1 and Rpn13 were the receptors-of-choice for the UBL domain (18). However, as a relatively easy to unfold substrate was used, it remained unclear to what extent the proteasome’s unfolding ability is affected by the targeting mode or receptor choice. By using substrates containing GFP variants of different stabilities with wild-type and receptor-mutant proteasomes, and using gel-based assays that can detect partial degradation, we show here that UBL-targeted substrates are degraded more efficiently than Ub_4_-targeted substrates, and that this difference seems to be correlated with reliance on the Rpn13 receptor.

## Results

### Superfolder GFP (sGFP) is not degraded by the proteasome when targeted via a linear Ub_4_ signal

We first examined the ability of the proteasome to degrade a substrate consisting of superfolder GFP (sGFP) with a linear Ub_4_ tag on the N-terminus and a C-terminal unstructured tail of 44 amino acids (including a terminal hexa-histidine tag), Ub_4_-sGFP-38-His_6_. The substrate was incubated with purified yeast proteasome, and the reaction mixture was run on an SDS-PAGE gel and visualized and quantified using the inherent fluorescence of GFP. There was essentially no degradation of sGFP as determined by quantifying the total fluorescence (**Figure 1A**; green). However, there was substantial “clipping” of the tail (**Figure 1A**; red and blue) as has been seen previously with other GFP-containing substrates (16, 20). Clipping could occur because the unstructured tail is too short to initiate processive unfolding or because after the proteasome initiates degradation, it is unable to unfold sGFP, stalls, and eventually releases the substrate. Longer initiation regions improve degradation of some proteasomal substrates (18, 26), but we found replacing the tail with an 108-amino acid long tail (Ub_4_-sGFP-102-His_6_) only modestly increased the rate and extent of complete degradation, and clipping remained the predominant outcome of degradation (**Figure 1B, E, F**). On the other hand, replacement of sGFP with a circular permutant of GFP (in which the 8^th^ ß-strand is the C-terminus instead of the 11^th^), cp8sGFP, led to rapid degradation of the substrate with very little clipping (**Figure 1C**), consistent with the hypothesis that the proteasome stalls while trying to unfold the highly stable sGFP.

**Figure 1.**
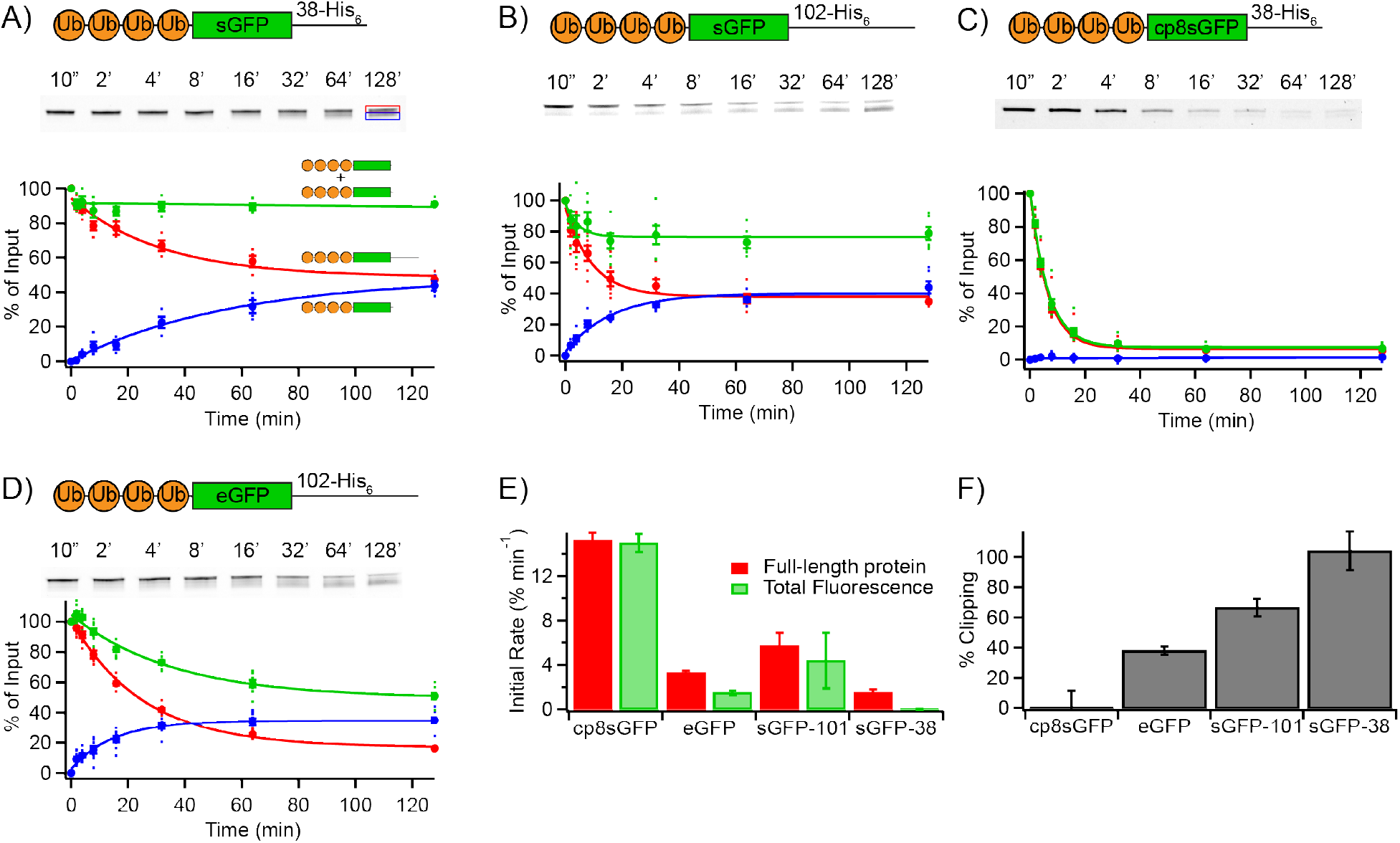
Degradation of Ub_4_-GFP substrates by the proteasome. **A**) Degradation of 100 nM Ub_4_-sGFP-38-His_6_ by 100 nM yeast proteasome. Representative gel shows disappearance of fulllength protein (red box) and appearance of clipped protein (blue box). The amounts of full-length protein (red circles), clipped protein fragment (blue circles), and total fluorescence (green circles) are shown as a percentage of the full-length substrate presented to the proteasome at the beginning of the reaction. Dots are results from individual experiments, and error bars represent the SEM of 4 experiments. Curves are global fits to single exponentials. **B**-**D**) As in A, but for Ub_4_-sGFP-102-His_6_, Ub_4_-cp8sGFP-38-His_6_, and Ub_4_-eGFP-102-His_6_. n = 6, 8 and 8. **E)** Initial rates of degradation from A-D for either disappearance of full-length protein (red) or disappearance of fluorescence (green). **F)** Percentage of full-length protein that was clipped rather than being completely degraded, as calculated by dividing the amplitude of clipped protein formation by the amplitude of disappearance of full-length protein.

Additional experiments support that the linear Ub_4_ tag is capable of targeting proteins to the proteasome, but the ability to successfully unfold and degrade the substrates depends on their stabilities. First, we confirmed that the Ub_4_-sGFP substrate did bind to the proteasome, as it was able to inhibit the degradation of a conventionally ubiquitinated substrate (**Supporting Figure S1**). As expected, clipping of the sGFP substrate and degradation of the cp8sGFP substrate were both greatly slowed by proteasome inhibitors (**Supporting Figure S2A-B**). However, the addition of free polyubiquitin chains slowed degradation of cp8sGFP but not clipping of sGFP, suggesting that clipping does not depend on ubiquitin (**Supporting Figure S2C-D**). We observed that clipping of sGFP could be mediated by either the 20S or 26S proteasome, as both free 20S core particle or doubly-capped 26S proteasome (reconstituted by the addition of excess purified 19S particle to purified 20S core particle) clipped the sGFP substrate similarly. In contrast, the cp8sGFP substrate was clipped by 20S proteasome but degraded once any 19S particle was added (**Supporting Figure S3**). Replacing sGFP with the less stable enhanced GFP (eGFP) led to intermediate levels of clipping and degradation (**Figure 1D, F**). In sum, while the linear Ub_4_ tag targets proteins to the proteasome, their unfolding and degradation depends on their stabilities. If substrates are too stable to be unfolded rapidly, they are instead clipped in a ubiquitin-independent manner, shortening and presumably inactivating the unstructured initiation site and thereby preventing re-targeting.

### Targeting via the UBL domain results in successful degradation of sGFP

Next, we replaced the Ub_4_ targeting sequence on sGFP with the UBL domain from yeast Rad23 (UBL-sGFP-102-His_6_). The proteasome was able to robustly degrade this substrate (as had been previously described (15)) with little clipping observed (**Figure 2A, E**). Degradation proceeded through a transient long fragment (blue curve), such that the rate of disappearance of full-length protein was much faster than the rate of disappearance of fluorescence (**Figure 2D**). A shorter persistent fragment (purple curve) was also produced, which presumably has an unstructured tail too short to serve as an initiation site. This fragment was likely produced directly from the full-length substrate, analogous to clipping of the Ub_4_-containing substrates via a proteasome-dependent but UBL-independent pathway (see **Figure 4** below). As with the Ub_4_-containing substrates, replacing sGFP with the less stable cp8sGFP (**Figure 2B**) reduced the difference between full-length and fluorescence degradation rates (**Figure 2D**) and reduced the extent of clipping (**Figure 2E**) such that the cp8sGFP substrate was degraded without the detection of any intermediates or formation of clipped fragment. (We note that this substrate also ran as a thicker band on a gel, potentially obscuring detection of intermediates). Only small differences were seen between eGFP and sGFP (**Figure 2C**). In all cases, the extent of complete degradation of a given GFP protein was larger with a UBL tag then with a Ub_4_ tag, indicating that the UBL domain is better able to promote the unfolding and degradation of the substrate by the proteasome.

**Figure 2.**
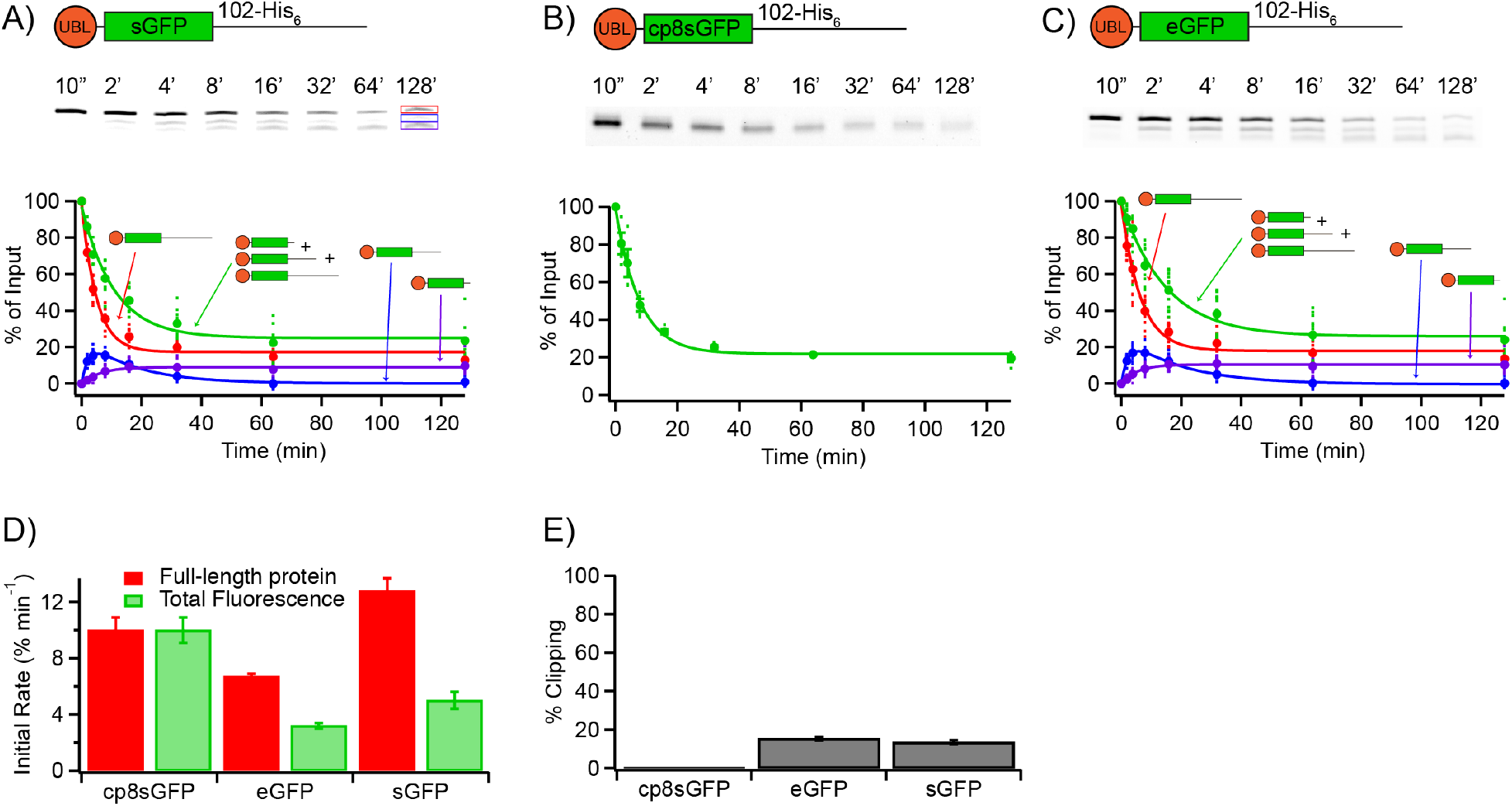
Degradation of UBL-GFP substrates by the proteasome. **A**) Degradation of 100 nM UBL-sGFP-102-His_6_ by 100 nM yeast proteasome. Representative gel shows disappearance of full-length protein (red box) and appearance of longer (blue box) and shorter (purple box) clipped protein. The amounts of full-length protein (red circles), longer partially degraded protein (blue circles), shorter clipped protein (purple circles), and total fluorescence (green circles) are shown as a percentage of the full-length substrate presented to the proteasome at the beginning of the reaction. Dots are results from individual experiments, and error bars represent the SEM of 8 experiments. Curves are global fits to single exponentials. **B**-**C**) As in A, but for UBL-cp8sGFP-102-His_6_ and UBL-eGFP-102-His. n = 14 and 4. In C, minimal clipped protein was formed, so only disappearance of total fluorescence is quantified. **D)** Initial rates of degradation from A-D for either disappearance of full-length protein (red) or disappearance of fluorescence (green). **E)** Percentage of full-length protein that was clipped (formed smaller fragment) rather than being degraded.

### Rates and extents of degradation are dependent on ATPase rate

It had previously been shown that for the Ub_4_-cp8sGFP-38-His_6_ substrate, slowing proteasomal ATP hydrolysis with ATP/ATPγS mixtures led to a proportional decrease in GFP degradation (17). The authors concluded from this result that unlike bacterial ATP-dependent proteases, the proteasome is highly processive and doesn’t release GFP sufficiently for refolding to occur. That is, even at low ATPase rates, *k*_refold_ << *k*_unfold2_ (**Figure 3A**). However, as we have shown, this circularly permuted substrate is much easier for the proteasome to unfold than eGFP or sGFP, and might indeed unfold catastrophically. We therefore set out to determine the effect of reducing ATP hydrolysis rates on unfolding and degradation of our substrates using ATPγS.

**Figure 3.**
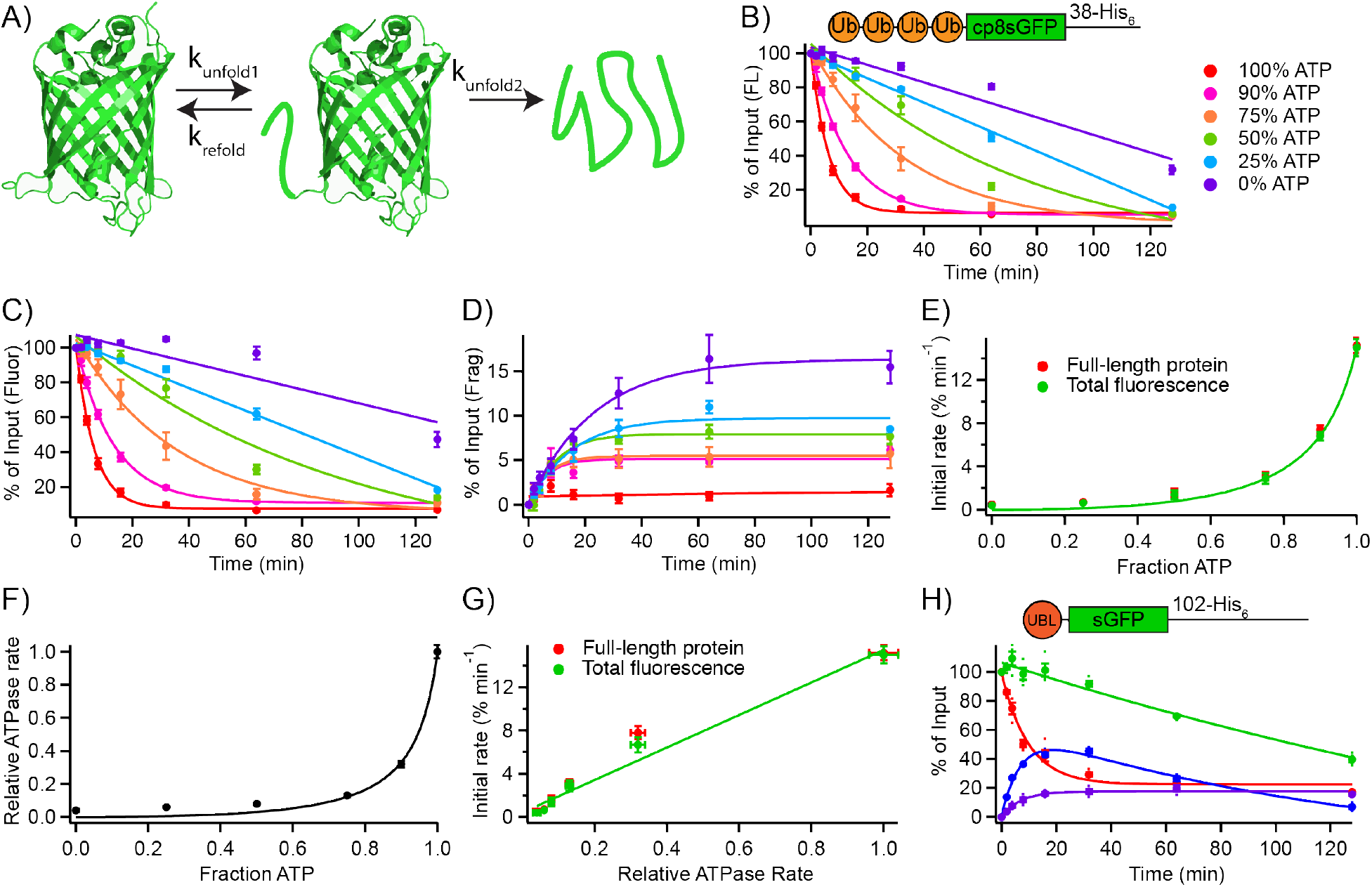
ATPγS slows degradation and ATPase activity similarly. **A**) Model for stepwise degradation of GFP, in which ATP-dependent extraction of the C-terminal ß-strand (*k*_unfold1_) may lead to a stable intermediate which can either refold (*k*_refold_) or be fully unfolded by additional pulls of the protesomal machinery (*k*_unfold2_). **B-D**) Effect of replacing ATP with ATPγS on disappearance of full-length Ub_4_-cp8sGFP-38-His_6_ (**B**), disappearance of fluorescence (**C**) and appearance of clipped GFP-containing fragment (**D**). Error bars represent the SEM of 4 experiments, except for 100% ATP (8). Conditions were as in **Figure 1B. E**) Initial rates from B and C as a function of fraction ATP. **F**) ATPase rates as a function of fraction ATP. Error bars represent the SEM of 4-10 experiments. **G**) Initial rates from B and C as a function of ATPase rates from F. **H**) Degradation of 100 nM UBL-sGFP-102-His_6_ by 100 nM yeast proteasome (as in **Figure 2A**) in the presence of 75% ATP/25% ATPγS. Error bars represent the SEM of 4 experiments.

In contrast to previous results (which were conducted using mammalian proteasome, not yeast proteasome), we found that replacing even a small fraction of ATP with ATPγS led to a precipitous drop in degradation rates for Ub_4_-cp8sGFP-38-His_6_ (**Figure 3B-D**). At higher ATPγS concentrations there was also a substantial increase in the extent of clipping (**Figure 3E**). This result was surprising, because we had not anticipated the formation of a stable unfolding intermediate with the cp8sGFP substrate. However, this analysis assumes that replacing ATP with ATPγS will linearly decrease the ATPase rate, which to our knowledge had not been tested. Using a coupled lactate dehydrogenase/pyruvate kinase ATPase assay, we found that even a small fraction of ATPγS is able to dramatically decrease the ATPase activity of the proteasome, consistent with a cooperative ATPase mechanism (**Figure 3F**). Similar results were seen using a malachite green assay (data not shown), indicating the results were not due to ATPγS inhibition of pyruvate kinase. Thus, there is indeed a linear dependence of degradation rate on ATPase rate for the cp8sGFP substrate (**Figure 3G**), consistent with a highly processive degradation mechanism for cp8sGFP, although important differences in terms of sensitivity to ATPγS are noted relative to mammalian proteasome. The increase in clipping seen at higher ATPγS concentrations indicates that slowing degradation sufficiently can cause the non-productive Ub-independent clipping pathway to become competitive with degradation even for this less stable substrate.

We next examined the degradation of the more stable sGFP-containing substrate, UBL-sGFP-102-His_6_. Replacing 25% of the ATP with ATPγS, which reduced the ATPase rate ~8-fold, reduced the initial rate of fluorescence loss by ~10-fold, about twice the effect seen with the cp8-sGFP substrate (an ~5-fold effect) (**Figure 3H** vs **Figure 2A**). Interestingly, the initial disappearance of full-length protein was only reduced by 1.6-fold, suggesting that formation of the longer clipped intermediate (*k*_unfold1_) is less sensitive to ATP concentration than unfolding of the rest of GFP (*k*_unfold2_). The extent of clipping was also doubled from ~10% to ~20%. Overall, these results indicate that although there is some ability for this more stable GFP to refold upon initial unfolding, even at low ATPase rates processive degradation is the predominate pathway taken. Thus, at least when targeted via the UBL domain, the proteasome does indeed processively degrade even a highly stable protein with little stalling. For comparison, ClpXP requires only an ~10% drop in ATPase rates for an ~90% decrease in degradation rates of sGFP (20).

### Partially degraded protein can be rebound and degraded

The larger GFP-containing fragment that appears and then decrease in intensity during degradation of UBL-targeted substrates (e.g., **Figure 2A**) could represent transiently-stalled protein bound to the proteasome before it has been degraded. Alternatively, it could indicate that partially degraded fragments can be released by the proteasome and then re-acquired via the N-terminal proteasome-binding tag (**Figure 4A)**. If those partially degraded fragments retain long enough initiation regions at the C-terminus of the protein, this would give the proteasome a second opportunity to attempt to unfold and degrade GFP (**Figure 4A**; dashed portion). To distinguish between these possibilities, we set up a pulse-chase experiment using the UBL-sGFP-102-His_6_ substrate in the presence of 25% ATPγS, thereby increasing the extent of transient fragment formation with a peak of fragment formation at ~9 minutes (**Figure 3H**). The addition of an excess of GST-UBL_Rad23_ at the peak of fragment formation will prevent (or greatly slow) rebinding of fully dissociated substrate to the proteasome via the UBL domain. If fragment is already associated with the proteasome, addition of competitor should not affect degradation of the fragment. However, if fragment dissociates and rebinds, its degradation should be slowed. We found that addition of the competitor slows the disappearance of the larger transient intermediate (**Figure 4B, C**, blue curve) and largely prevents further degradation of GFP (red and green). We conclude that the complete model of **Figure 4A** is most likely to hold, in which substrate is often released after initial engagement and partial degradation, and then is rebound and re-engaged by the proteasome. If the substrate is clipped more extensively, the shortened unstructured initiation site may no longer be able to productively engage with the proteasome. This more extensive clipping, analogous to the clipping we observed of Ub_4_-substrates, does not depend on the UBL domain, as it continued (and in fact increased) upon addition of GST-UBL_Rad23_, and addition of the competitor at the beginning of the assay slowed formation of the larger fragment and led to increased formation of the smaller fragment (**Figure 4B, D**). Indeed, kinetic modeling to the complete model of **Figure 4A** was able to simultaneously fit all three data sets (**Supporting Figure S4**). These results are also consistent with the ability of ATPγS to increase clipping (**Figure 3D, H**); an increase away from unfolding would increase clipping and thus the formation of no-longer engageable substrate fragments. Indeed, our kinetic model is able to reasonably fit the degradation of UBL-sGFP-102-His_6_ in the absence of ATPγS if 25% ATPγS slows the unfolding rate constant by ~5-fold and doesn’t affect any other rate constants (**Supporting Figure S4**). The ability to repeatedly release and rebind the substrate when degradation stalls might make it easier to degrade physiological substrates where the proteasome-binding element is distal to the initiation region (as opposed to many proteins where ubiquitination occurs on the initiation region, such that ubiquitin on the substrate would be removed prior to fragment release), and the ability of the proteasome to trim unstructured initiation sites may serve as a counterbalance, preventing the same substrate from being repeatedly fruitlessly engaged.

**Figure 4.**
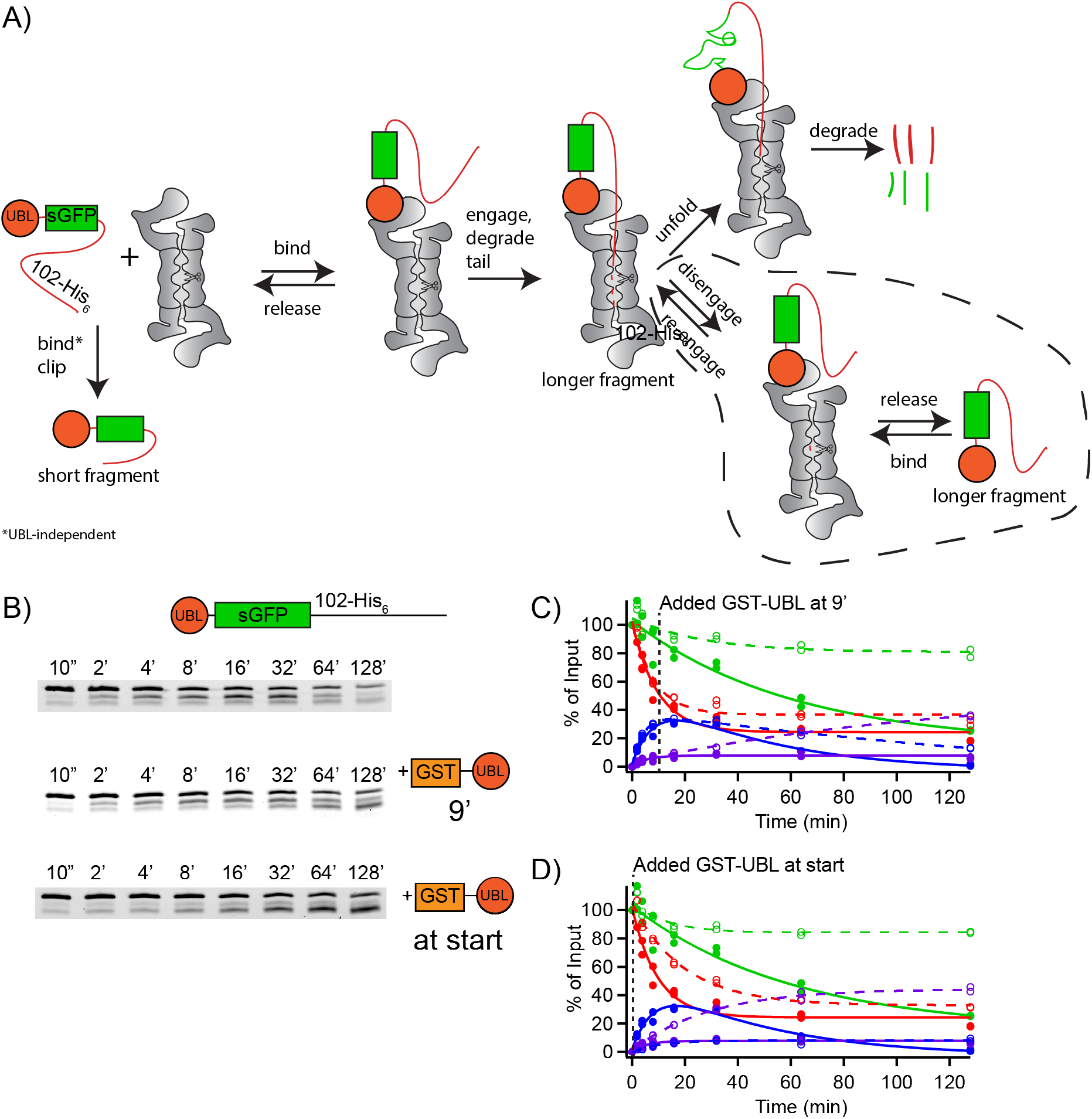
Partially degraded protein can be rebound and degraded. **A**) Model for degradation of UBL-sGFP-102-His_6_ substrate. Substrate can be clipped, forming a short fragment, in a UBL-independent (but proteasome-dependent) process, or can be bound via the UBL, engaged, the tail is partly degraded to produce the longer fragment which can either be exclusively unfolded and completely degraded (top pathway) or can alternatively be released and rebound repeatedly until it is degraded. **B**) Representative gels for degradation assays with 100 nM proteasome and 100 nM UBL-sGFP-102-His_6_ in which 5 μM GST-UBL_Rad23_ (or buffer) is added after 9 minutes or at the start of the assay. **C, D**) Quantification of pulse-chase assays from B as in **Figure 2A**. Closed symbols are mock addition and open symbols are with addition of GST-UBL_Rad23_. Dashed line shows approximate time of addition.

### Rpn13 is primarily responsible for strong unfolding with UBL targeting

Ub_4_ binds ~4 times more weakly to the proteasome than UBL (15, 19). However, decreasing the affinity 4-fold in our model has only a minor impact on simulated degradation, such that the proteasome should still largely degrade the substrate with minimal clipping. Given that a cp8sGFP substrate is degraded easily by either proteasome, there are likely also stability-linked differences in the engagement and unfolding rates. Ub_4_ and UBL substrates might be positioned differently at the proteasome, affecting engagement and perhaps even unfolding ability. We were therefore curious to know if one or more of the intrinsic proteasomal ubiquitin receptors were responsible for the proteasome’s ability to degrade hard-to-unfold substrates with UBL but not Ub_4_ targeting. The Matouschek lab had previously shown that the linear Ub_4_ degron (or, more specifically, the Ub_4_-cp8sGFP-38-His_6_ substrate) is recognized and degraded by both Rpn10 and Rpn13 (but not Rpn1), while the corresponding UBL substrate is degraded even if all three receptors are removed, but requires Rpn1 or Rpn13 for maximal degradation efficiency (18). To determine the requirements for a more difficult to unfold substrate, we used Ub_4_- or UBL-eGFP-102-His_6_, which is degraded very efficiently with the UBL signal, and with moderate efficiency with the Ub_4_ signal (**Figure 1D, Figure 2B**). When targeted via the Ub_4_ signal, individual proteasome receptor mutants (Rpn1ΔT1, Rpn10ΔUIM and Rpn13-pru, each of which use point mutations to disrupt ubiquitin binding (25)) had modest effects, generally increasing the amount of clipping and, in the case of Rpn13-pru, reducing the rate of overall degradation (disappearance of fluorescence) (**Figure 5A, B; Supporting Figure S5**). Double mutants retaining only one receptor (Rpn1-only, Rpn10-only and Rpn13-only) again had increased clipping but only small effects on degradation rates, suggesting that all of the receptors are capable of mediating at least some degradation of this substrate, although none of the individual receptors are able to do so particularly efficiently. In contrast, targeting via the UBL domain was much more receptor-specific. Mutating either Rpn1 or Rpn13 modestly increased clipping and, in the case of Rpn13-pru, reduced overall degradation rates (**Figure 5C, D**). Surprisingly, mutating Rpn10 actually increased overall degradation rates, suggesting that binding of the UBL domain to Rpn10 might be non- or counter-productive. The effects of single-receptor-containing proteasomes were even more striking. Rpn13-only proteasome behaved essentially like wildtype, while Rpn1-only and Rpn10-only proteasome were almost incapable of unfolding and degrading GFP, with greatly reduced overall degradation rates and greatly elevated clipping. Indeed, these mutants essentially convert the well-degraded UBL substrate into the poorly-degraded Ub_4_ substrate, indicating that Rpn13 is primarily responsible for the enhanced unfolding and degradation of UBL-containing substrates, although the combination of Rpn10 and Rpn1 is able to partially compensate for the loss of Rpn13’s ability to bind the UBL domain. We confirmed that the inability to degrade was not simply due to a lack of substrate binding, as the UBL-cp8sGFP-102-His_6_ substrate was still degraded by the Rpn1-only proteasome with an initial rate only 2-3-fold reduced from WT without any evidence of fragment formation (**Supporting Figure S6**).

**Figure 5.**
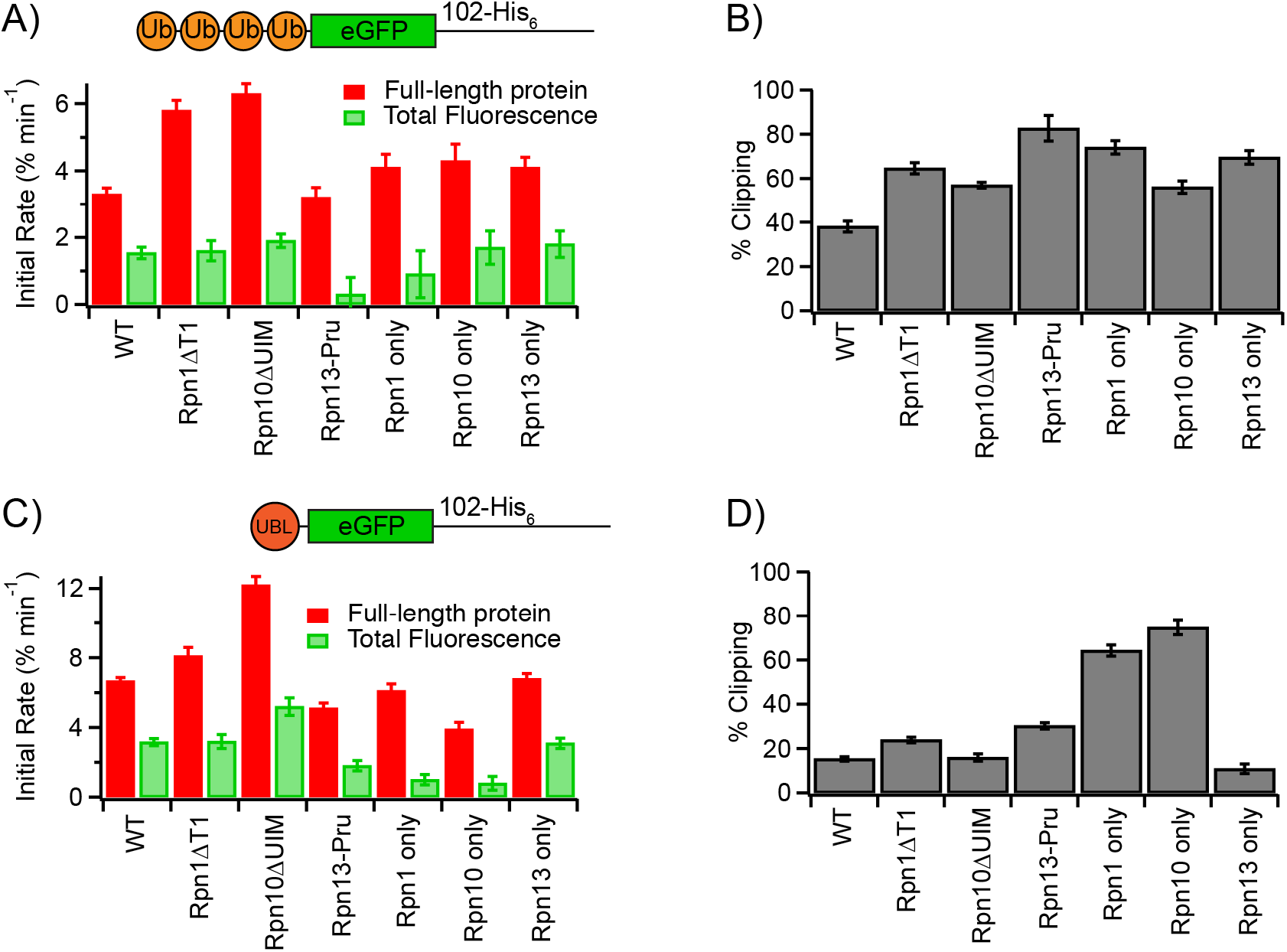
Effects of proteasome receptor mutations on degradation of eGFP-containing substrates. **A**) Initial rates of degradation for Ub_4_-eGFP-102-His_6_ for either disappearance of fulllength protein (red) or disappearance of fluorescence (green). Error bars are SEM derived from curve-fitting 4 replicate experiments. **B**) Percentage of full-length Ub_4_-eGFP-102-His_6_ that was clipped rather than being completely degraded. **C,D**) As in A and B, but for UBL-eGFP-102-His_6_. Error bars are SEM derived from curve-fitting 4 replicate experiments.

## Discussion

GFP is commonly used as a model substrate to study protein degradation. Here we show that both the ubiquitin binding tag (Ub_4_ vs UBL) and the GFP variant’s stability are critically important for determining whether GFP is successfully unfolded and degraded or whether degradation is terminated unsuccessfully. UBL-targeted substrates with the same stability are degraded more easily than Ub_4_-targeted substrates. This extra unfolding ability is mediated, at least in part, by the Rpn13 ubiquitin receptor. We had previously proposed that binding of ubiquitinated proteins to Rpn13 might help activate the proteasome’s unfolding ability by shifting the conformation from the s1 substrate-binding state to s3-like substrate processing states; in the kinetic model of **Figure 4A** this would correspond to increasing the initial engagement, degradation, and/or unfolding rates (16, 25). The same hypothesis could explain the UBL versus Ub_4_ results we see, given that Ub_4_ substrates rely primarily on Rpn10, while UBL substrates rely primarily on Rpn13 (18). Consistent with this hypothesis, release of polyubiquitin chains from Rpn13 has been suggested to slow multiple turnover degradation by the proteasome, suggesting engagement of Rpn13 may keep the proteasome from returning to the s1 state for additional rounds of substrate binding (18). An alternative possibility is that differences between Ub_4_ and UBL stem from the need to pull the proteasome-binding tag away from its receptor during translocation. Extracting a single UBL domain may be easier than extracting Ub_4_, which might more easily re-bind during the process. However, the UBL domain binds ~4-fold more tightly to the proteasome than Ub_4_ (15, 19), arguing against this model.

It has long been puzzling why so many ubiquitin receptors and shuttle proteins were needed for proteasomal function. Our results suggest that shuttle proteins like Rad23 may be used to bring harder-to-unfold proteins to the proteasome by tethering them to Rpn13 and activating an enhanced unfolding ability, while easier-to-unfold proteins are able to be degraded without this extra assistance by directly docking to Rpn10. Recently, it has been suggested that there is at least one additional ubiquitin or UBL receptor on the proteasome, as even a triple-receptor mutant can still degrade a UBL-cp8sGFP-102-His_5_ substrate (18). Our results indicate that if indeed an additional ubiquitin or UBL receptor exists, it does not support unfolding of difficult to unfold substrates such as non-circularly permuted GFP.

Our results also indicate that UPS substrates where the two-component degron (proteasome binding tag and unstructured initiation region) is split between different portions of the protein may be degraded more robustly than those where the initiation region and binding tag are near one another in the primary sequence. Separation allows partially degraded protein fragments to be re-targeted to the proteasome, such that a lower processivity is required in order to get complete degradation of the substrate. However, this is balanced by the ability of the proteasome to trim unstructured initiation regions in a binding-tag independent fashion (**Figure 4A**), which may prevent futile cycles of attempted degradation. Indeed, “masked” initiation sites, which are only exposed when degradation is needed (30), may be important both for preventing premature degradation and for preventing premature destruction of the initiation site.

Finally, these results have implications for the use of GFP as a tool in assays and screens that measure the global health of the UPS or that examine the fate of individual proteins. Given GFP’s high stability, it seems advisable to use a circular permutant in cases where one simply wants to know if an attached protein degrades, or how robust the UPS is for degradation of easy-to-unfold proteins. A more stable GFP such as sGFP could instead be used to look at the ability of the UPS to degrade more challenging substrates, and again, the nature of the targeting mechanism may play a role in determining how robust of a proteasome response is required for degradation.

## Experimental Procedures

### Constructs

Plasmids expressing Ub_4_-sGFP-38-His_6_ in pGEM3Zf+, Ub_4_-cp8sGFP-38-His_6_ in pGEM3ZF+ and UBL-sGFP-102-His_6_ in pETDuet were kind gifts from the Matouschek lab. A plasmid expressing Ub_4_-sGFP-102-His_6_ was made by replacing the UBL domain in UBL-sGFP-102-His_6_ with a Ub_4_ domain by restriction cloning. A plasmid expressing UBL-CP8sGFP-102-His_6_ was made by replacing the sGFP from UBL-sGFP-102-His_6_ with the CP8sGFP from Ub_4_-CP8sGFP-38-His_6_ by restriction cloning. Plasmids expressing eGFP substrates were made by replacing sGFP with eGFP by restriction cloning. A plasmid expressing GST-UBL_Rad23_ was made by inserting UBL_Rad23_ from yeast into pDEST15. Plasmid sequences are available upon request.

### Proteasome Purification

Yeast (*S. cerevisiae*) proteasome was purified from wild type strain YYS40 or receptor mutant strains using a 3X-FLAG-tagged copy of Rpn11 subunit of the 19S particle as described previously (16, 25). Cells were grown overnight in YEPD or selective media before being transferred to YEPD media for large scale growth at 30°C. Cells were harvested by centrifugation and resuspended in buffer containing 25 mM Tris pH 7.5, 10 mM MgCl_2_, 4 mM ATP, 1 mM DTT, 10% (v/v) glycerol, and a 2X ATP-regeneration system (ARS) consisting of 0.5 mg/mL creatine phosphokinase and 20 mM phosphocreatine. Cells were then lysed by homogenization and the lysate was pH adjusted to 7.5 using 1 M Tris base before high-speed centrifugation. The supernatant was supplemented with 5 mM ATP and 1X ARS, filtered, and incubated with anti-FLAG M2 Affinity Gel at 4°C while rotating for 2 hours. The resin was washed 3 times with 25 mM Tris pH 7.5, 5 mM MgCl_2_, 2 mM ATP, 1 mM DTT, and 10% (v/v) glycerol. The proteasome was eluted with 100 μg/mL 3X Flag peptide. Peak elutions were pooled, and concentration was determined by Bradford assay. Proteasome was flash frozen and stored at −80 °C until use. To purify proteasome in the presence of ATPγS, the final wash and the elution buffer contained 2 mM ATPγS instead of ATP. ATPγS was purchased from Cayman Chemical, dissolved in ddH2O and 1 M Tris-Cl to a pH of ~8, and stored as 10 mM ATPγS at −80°C until use. To purify 19S particle, resin was washed 3 times with 25 mM Tris pH 7.5, 500 mM NaCl, 5 mM MgCl_2_, 2 mM ATP, 1 mM DTT, and 10% (v/v) glycerol and then 1 time with 25 mM Tris pH 7.5, 5 mM MgCl_2_, 2 mM ATP, 1 mM DTT, and 10% (v/v) glycerol before elution. To purify 20S particle, strain yMDC11, containing a 3X-FLAG-tagged copy of Pre1, was used, ATP, ARS and MgCl_2_ were omitted from buffers, and resin was washed 3 times with 25 mM Tris pH 7.5, 500 mM NaCl, 1 mM DTT, and 10% (v/v) glycerol and then 1 time with 25 mM Tris pH 7.5, 1 mM DTT, and 10% (v/v) glycerol before elution.

### Bacterial protein overexpression and purification

Ub_4_-sGFP-38-His_6_ and Ub_4_-cp8sGFP-38-His_6_ were transformed into Rosetta(DE3)pLysS cells, grown in LB media at 37°C to an OD_600_ of 0.6, induced with 0.4 mM IPTG at an OD_600_ of 0.6, and grown for 12-18 hours. Cells were spun down, resuspended in lysis buffer (50 mM sodium phosphate, 300 mM NaCl, 10 mM imidazole, pH 8.0), lysed by high-pressure homogenization, bound to a NiNTA column, washed, and eluted with increasing concentrations of imidazole. The purest fluorescent fractions were pooled (as determined by SDS-PAGE), and the concentration was determined by absorbance at 280 nm. Proteins were flash frozen and stored at −80°C.

UBL-eGFP-102-His_6_ and Ub_4_-eGFP-102-His_6_ were purified identically, but they were transformed into BL21(DE3) cells and overexpressed in LB media at 30° and 37°C, respectively.

UBL-CP8sGFP-102-His_6_ and Ub_4_-sGFP-102-His_6_ were purified identically, but they were transformed into BL21 (DE3) cells and overexpressed in autoinduction media at 37°C (31).

GST-UBL_Rad23_ was transformed into BL21AI cells, grown in LB media at 37 °C to an OD_600_ of 0.6, and induced with 0.1% arabinose for 3 hours. Cells were spun down, resuspended in PBS, lysed by high-pressure homogenization, and loaded onto a Glutathione-agarose column. After washing with PBS, protein was eluted with GSH elution buffer (100 mM Tris-Cl, 100 mM NaCl, 10 mM GSH, 1 mM DTT pH 8.0) and then dialyzed against 25 mM HEPES, 5 mM MgCl_2_, 1 mM DTT, 10% (v/v) glycerol pH 7.4. Concentration was determined using a Pierce 660 assay. Proteins were flash frozen and stored at −80°C.

Neh2Dual-Barnase-DHFR substrate was expressed, purified, labeled and ubiquitinated as described previously (25). K63-linked chains were enzymatically synthesized and purified as described previously (16).

### Degradation Assays

Degradation assays were conducted using 100 nM proteasome and 100 nM fluorescent substrate over a 2-hour time course. Reactions were carried out at 30° C in degradation buffer (50 mM TrisCl, 5 mM MgCl_2_, 5% (v/v) glycerol, 1 mM ATP, 10 mM creatine phosphate, 0.1 mg/mL creatine kinase, and 1% DMSO, pH 7.5). Reactions contained 1 mg/mL BSA to prevent nonspecific loss of fluorescence. For reconstitution experiments, 20S core particle was preincubated with 19S core particle in the presence of degradation buffer for 10 minutes at 30 °C before beginning the assay by the addition of substrate plus BSA. At designated time points, samples were removed and placed into SDS-PAGE loading buffer to quench the reaction; samples were not heated to prevent denaturation of GFP. SDS-PAGE gels were analyzed by fluorescence imaging on a Typhoon FLA 9500, and the resulting gel files were analyzed using ImageQuant (GE).

### ATPase Assays

Proteasome ATPase activity was measured using a coupled pyruvate kinase/lactate dehydrogenase assay in saturating ATP, which can be spectrophotometrically detected at 340 nm. Reactions contained 20 nM proteasome, 6.8 U/mL pyruvate kinase, 9.9 U/mL lactate dehydrogenase, 0.4 mM NADH, 2 mM phosphoenolpyruvate, 0.5 mM DTT, and various ATP/ATPγS ratios totaling 0.5 mM in a buffer consisting of 50 mM HEPES-KOH pH 7.5, 50 mM KCl, and 5 mM MgCl_2_. Reaction was run at 30°C in a 384 well plate with time points taken every 20 seconds for 20 minutes by a BioRad Benchmark Plus UV-Vis platereader.

### Native Gels

Proteasome was reconstituted as for degradation assays, then run on a 3.5% native gel and visualized using Suc-LLVY-AMC in the presence of 0.02% SDS (32).

### Kinetic Modeling

Kinetic modeling was carried out using COPASI software (33).

## Data availability

All data not in the manuscript will be shared upon request to DAK

## Acknowledgements

The authors thank Andreas Matouschek for sharing Ub_4_-cp8sGFP-38-His_6_, Ub_4_-sGFP-38-His_6_, and UBL-sGFP-102-His_6_ plasmids. The authors thank Aimee Eggler, David Smith and members of the Kraut lab for helpful discussions and feedback.

## Abbreviations

UPS: ubiquitin-proteasome system
GFP: green fluorescence protein
UFD: ubiquitin-fusion degradation
UBL: ubiquitin-like domain
sGFP: superfolder GFP

## References

1. Trauth, J., Scheffer, J., Hasenjäger, S., Biophysics, C. T. A., 2020 (2020) Strategies to investigate protein turnover with fluorescent protein reporters in eukaryotic organisms. AIMS Biophysics. 7, 90–118

2. Dantuma, N. P., Lindsten, K., Glas, R., Jellne, M., and Masucci, M. G. (2000) Short-lived green fluorescent proteins for quantifying ubiquitin/proteasome-dependent proteolysis in living cells. Nature Biotechnology. 18, 538–543

3. Heessen, S., Dantuma, N. P., Tessarz, P., Jellne, M., and Masucci, M. G. (2003) Inhibition of ubiquitin/proteasome-dependent proteolysis in Saccharomyces cerevisiae by a Gly-Ala repeat. FEBS Lett. 555, 397–404

4. Holmberg, C. I., Staniszewski, K. E., Mensah, K. N., Matouschek, A., and Morimoto, R. I. (2004) Inefficient degradation of truncated polyglutamine proteins by the proteasome. EMBO J. 23, 4307–4318

5. Maynard, C. J., Böttcher, C., Ortega, Z., Smith, R., Florea, B. I., Díaz-Hernández, M., Brundin, P., Overkleeft, H. S., Li, J.-Y., Lucas, J. J., and Dantuma, N. P. (2009) Accumulation of ubiquitin conjugates in a polyglutamine disease model occurs without global ubiquitin/proteasome system impairment. Proc Natl Acad Sci USA. 106, 13986–13991

6. McKinnon, C., Goold, R., André, R., Devoy, A., Ortega, Z., Moonga, J., Linehan, J. M., Brandner, S., Lucas, J. J., Collinge, J., and Tabrizi, S. J. (2016) Prion-mediated neurodegeneration is associated with early impairment of the ubiquitin-proteasome system. Acta Neuropathol. 131, 411–425

7. Thibaudeau, T. A., Anderson, R. T., and Smith, D. M. (2018) A common mechanism of proteasome impairment by neurodegenerative disease-associated oligomers. Nat Commun. 9, 1097

8. Moura, D. M. N., de Melo Neto, O. P., and Carrington, M. (2019) A new reporter cell line for studies with proteasome inhibitors in Trypanosoma brucei. Mol Biochem Parasitol. 227, 15–18

9. Bence, N. F., Sampat, R. M., and Kopito, R. R. (2001) Impairment of the ubiquitin-proteasome system by protein aggregation. Science. 292, 1552–1555

10. Greussing, R., Unterluggauer, H., Koziel, R., Maier, A. B., and Jansen-Dürr, P. (2012) Monitoring of ubiquitin-proteasome activity in living cells using a Degron (dgn)-destabilized green fluorescent protein (GFP)-based reporter protein. J Vis Exp. 10.3791/3327

11. Arquier, N., Vigne, P., Duplan, E., Hsu, T., Therond, P. P., Frelin, C., and D’Angelo, G. (2006) Analysis of the hypoxia-sensing pathway in Drosophila melanogaster. Biochem J. 393, 471–480

12. Khmelinskii, A., Keller, P. J., Bartosik, A., Meurer, M., Barry, J. D., Mardin, B. R., Kaufmann, A., Trautmann, S., Wachsmuth, M., Pereira, G., Huber, W., Schiebel, E., and Knop, M. (2012) Tandem fluorescent protein timers for in vivo analysis of protein dynamics. Nature Biotechnology. 30, 708–714

13. Wang, S., Tang, N. H., Lara-Gonzalez, P., Zhao, Z., Cheerambathur, D. K., Prevo, B., Chisholm, A. D., Desai, A., and Oegema, K. (2017) A toolkit for GFP-mediated tissue-specific protein degradation in C. elegans. Development. 144, 2694–2701

14. Bashore, C., Dambacher, C. M., Goodall, E. A., Matyskiela, M. E., Lander, G. C., and Martin, A. (2015) Ubp6 deubiquitinase controls conformational dynamics and substrate degradation of the 26S proteasome. Nat Struct Mol Biol. 22, 712–719

15. Yu, H., Kago, G., Yellman, C. M., and Matouschek, A. (2016) Ubiquitin-like domains can target to the proteasome but proteolysis requires a disordered region. EMBO J. 10.15252/embj.201593147

16. Reichard, E. L., Chirico, G. G., Dewey, W. J., Nassif, N. D., Bard, K. E., Millas, N. E., and Kraut, D. A. (2016) Substrate Ubiquitination Controls the Unfolding Ability of the Proteasome. J. Biol. Chem. 291, 18547–18561

17. Snoberger, A., Anderson, R. T., and Smith, D. M. (2017) The Proteasomal ATPases Use a Slow but Highly Processive Strategy to Unfold Proteins. Front Mol Biosci. 4, 18

18. Martinez-Fonts, K., Davis, C., Tomita, T., Elsasser, S., Nager, A. R., Shi, Y., Finley, D., and Matouschek, A. (2020) The proteasome 19S cap and its ubiquitin receptors provide a versatile recognition platform for substrates. Nat Commun. 11, 477

19. Singh Gautam, A. K., Martinez-Fonts, K., and Matouschek, A. (2018) Scalable In Vitro Proteasome Activity Assay. Methods Mol Biol. 1844, 321–341

20. Nager, A. R., Baker, T. A., and Sauer, R. T. (2011) Stepwise Unfolding of a β Barrel Protein by the AAA+ ClpXP Protease. J Mol Biol. 413, 4–16

21. Koodathingal, P., Jaffe, N. E., Kraut, D. A., Prakash, S., Fishbain, S., Herman, C., and Matouschek, A. (2009) ATP-dependent proteases differ substantially in their ability to unfold globular proteins. 284, 18674–18684

22. Dai, R. M., and Li, C. C. (2001) Valosin-containing protein is a multi-ubiquitin chain-targeting factor required in ubiquitin-proteasome degradation. Nat Cell Biol. 3, 740–744

23. Beskow, A., Grimberg, K. B., Bott, L. C., Salomons, F. A., Dantuma, N. P., and Young, P. (2009) A conserved unfoldase activity for the p97 AAA-ATPase in proteasomal degradation. J Mol Biol. 394, 732–746

24. Olszewski, M. M., Williams, C., Dong, K. C., and Martin, A. (2019) The Cdc48 unfoldase prepares well-folded protein substrates for degradation by the 26S proteasome. Commun Biol. 2, 29

25. Cundiff, M. D., Hurley, C. M., Wong, J. D., Boscia, J. A., Bashyal, A., Rosenberg, J., Reichard, E. L., Nassif, N. D., Brodbelt, J. S., and Kraut, D. A. (2019) Ubiquitin receptors are required for substrate-mediated activation of the proteasome’s unfolding ability. Sci Rep. 9, 14506

26. Inobe, T., Fishbain, S., Prakash, S., and Matouschek, A. (2011) Defining the geometry of the two-component proteasome degron. Nat Chem Biol. 7, 161–167

27. Shi, Y., Chen, X., Elsasser, S., Stocks, B. B., Tian, G., Lee, B.-H., Shi, Y., Zhang, N., de Poot, S. A. H., Tuebing, F., Sun, S., Vannoy, J., Tarasov, S. G., Engen, J. R., Finley, D., and Walters, K. J. (2016) Rpn1 provides adjacent receptor sites for substrate binding and deubiquitination by the proteasome. Science. 351, aad9421–aad9421

28. Grice, G. L., and Nathan, J. A. (2016) The recognition of ubiquitinated proteins by the proteasome. Cell. Mol. Life Sci. 73, 3497–3506

29. Lu, Y., Lee, B.-H., King, R. W., Finley, D., and Kirschner, M. W. (2015) Substrate degradation by the proteasome: a single-molecule kinetic analysis. Science. 348, 1250834

30. Tomita, T., Huibregtse, J. M., and Matouschek, A. (2020) A masked initiation region in retinoblastoma protein regulates its proteasomal degradation. Nat Commun. 11, 2019

31. Studier, F. W. (2005) Protein production by auto-induction in high-density shaking cultures. Protein Expr Purif. 41, 207–234

32. Elsasser, S., Schmidt, M., and Finley, D. (2005) Characterization of the Proteasome Using Native Gel Electrophoresis, Methods in Enzymology, Elsevier, 398, 353–363

33. Hoops, S., Sahle, S., Gauges, R., Lee, C., Pahle, J., Simus, N., Singhal, M., Xu, L., Mendes, P., and Kummer, U. (2006) COPASI--a COmplex PAthway Simulator. Bioinformatics. 22, 3067–3074

